# Analysing variations in pelage patterning among European *Felis silvestris silvestris* populations

**DOI:** 10.1101/2021.02.22.432262

**Authors:** David Reher, Clara Stefen

## Abstract

The pelage pattern of wildcats from six regions within the distribution range of the species is analysed to test the hypothesis that clear differences between different regions exist. In total, 98 furs were used, but with different distribution from the Eifel (Germany), Harz Mountains (Germany), Caucasus, Western Spain, Switzerland/France and Greece. Specimens were at least seven months old, sex was not considered. The characteristics used were adapted from the literature and included typical wildcat features like the tail and stripes on the neck. Polychoric correlations, as a binary version of Spearman’s rank correlation coefficient, were used as a measure of correlation between pelage characteristics. Pairwise tetrachoric correlations were computed with R 3.1.2., cluster analyses were conducted with SPSS 14.

No clear relationship between the pelage characteristics and the geographic distribution of the six studied wildcat populations were found, the hypothesis needs to be rejected. It is however suggested, that coat patterning of Caucasian wildcats to some degree differ from other European wildcat. The results, however, do not clearly corroborate this hypothesis as Caucasian wildcats were not grouped into closed clusters in all the cluster analyses. To a lesser extent, pelage characteristic differences may also exist between wildcats from Western Spain as well as South-Western Greece and other European regions, but the samples are too small to draw conclusions. To test these refined hypotheses, more specimens from the Caucasus, Western Spain and South-Western Greece (but also Switzerland/North-Eastern France) need to be collected and compared according to our protocol.

## Introduction

The wildcat *Felis silvestris* has a large distribution range which includes Western Europe, a large part of Africa, Arabia, South-West Asia and China. The species can roughly be divided into three subgroups in terms of geographical distribution: i.e. African (*F. silvestris lybica*), Asian (*F. silvestris ornata*) and European (*F. silvestris silvestris*) wildcats. These subspecies also show fairly different ecological niches and morphologies and therefore are referred to as morphotypes (Kitchener et al. 2009). Even though the existence of distinct *F. silvestris* subgroups is generally accepted, the systematic classifications of several *Felis* taxa as well as the number of subspecies of *F. silvestris* are under constant debate (Kitchener et al. 2009). More recently these are considered as separate species, which leaves a smaller geographic range for Felis silvestris with the subspecies F. s. silvestris from (Central) Europe, but now including Scotland, Sicily and Crete, F. s. grampia originally from Scotland but probably extinct and F. s. caucasica from the Caucasus and Turkey (Kitchener et al. 2017).

The question whether *F. silvestris* should be considered a polytypical species with a large distribution range or needs to be divided into several species thus has been decided, but more more aspects of taxon differentiation might be helpful still. There is a long-standing debate what defines a wildcat and how to distinguish between different wildcat forms and domestic cats (i.e. Weigelt 1961; Ragni and Possenti 1996; Yamaguchi et al. 2004;). Substantial literature exists on the differentiation of the European wildcat and domestic cats considering several characteristics that were shown either to be diagnostic or, otherwise, not reliable (i.e. Piechocki 1990; Beaumont et al. 2001; Kitchener et al. 2005; Müller 2011). The most prominent and generally accepted characteristics are cranial volume and cranial index as well as intestine length and intestine index (Schauenberg 1969, 1971, 1977) and the thick and blunt tail with distinct black rings. Kitchener et al. (2005) identified seven pelage characteristics that most significantly differentiate wildcats, domestic cats and hybrids in Scotland, and Krüger et al. (2009) argue for five characteristics to distinguish these cat groups in Thuringia (Germany).

To differentiate taxa knowledge about the variation into metric and morphological traits is necessary. For the wildcat, particularly the variation of craniometric features has been described including changes with ontogeny (i.e. Kratochvíl and Kratochvíl 1970; Kratochvíl 1973; Schauenberg 1971b; Piechocki 1990; Stefen and Heidecke 2011, 2012). Less is known concerning the variability or even regional differences of pelage patterns and characteristics. Eckstein (1919) probably was the first to describe the variation in the stripes on neck and back and of tail shape and pattern in European wildcats. For wildcats from Germany, Vogt (1985) illustrated the variation in tail shape and pattern in Rhineland-Palatinate, and Müller (2011) depicted the overall variation of several pelage characteristics in Hessia. Randi and Possenti (1996) found that wildcats, domestic cats and Sardinian cats can well be distinguished on the basis of fur characteristics.

The morphotype *F. silvestris silvestris* is a more forest-dwelling variant than the others and predominant in European regions (Kitchener et al. 2009). However, its distribution throughout Europe is not coherent (Fig. 1) which allows the question whether morphological differences exist between geographically separated populations. For detailed analyses of feline coats, somatic regions of the cat (Fig. 2) and key pelage properties (Fig. 3) have been defined. It is known that between *F. silvestris* subgroups these characteristics differ to some degree. Accordingly, coat properties allow for discrimination between *F. silvestris* morphotypes (see Ragni and Possenti 1996 for differentiation of *F. silvestris catus, lybica* and *silvestris*). Kitchener et al. (2005) came to similar results in their analysis of differences between Scottish wildcats, hybrids and domestic cats by including skull parameters and intestinal length, which are known to be reliable discrimination parameters. They also specified coat properties best suited for differentiation: (1) extent of dorsal stripe, (2) shape of tail tip, (3) distinctness of tail bands, (4) presence/absence of broken stripes and (5) spots on flanks and hindquarters, (6) shape and number of stripes on nape and (7) shoulders (Fig. 3).

**Fig. 1.**
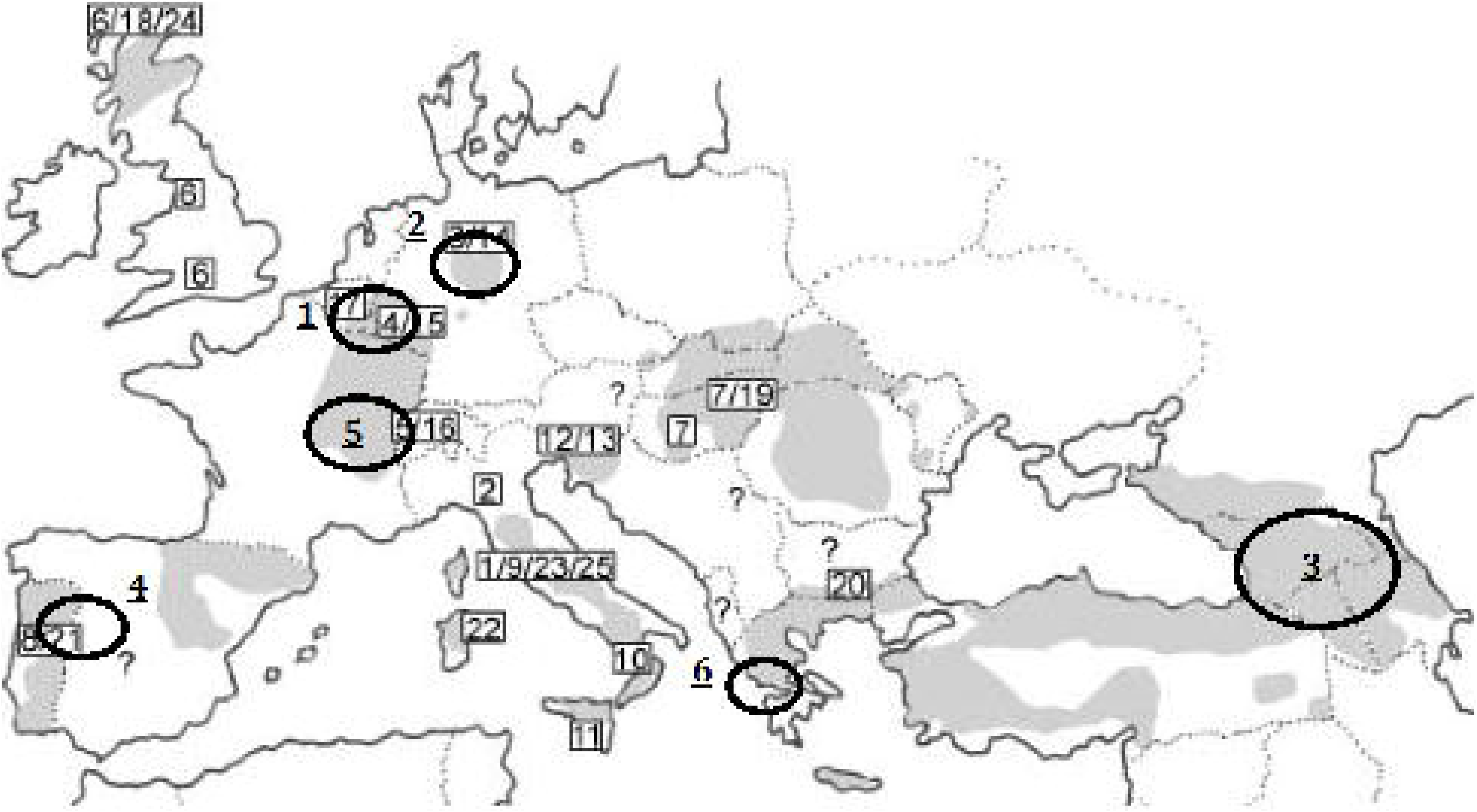
Distribution of *Felis silvestris silvestris* in Europe. Here, individuals approximately found in the circled areas were analysed. 1: Eifel (n=21), 2: Harz (n=49), 3: Caucasus (n=12), 04: Western Spain (n=4), 05: Switzerland/North-Eastern France (n=10), 06: South-Western Greece (n=2) (adapted from Pierpaoli et al., 2003).

**Fig. 2.**
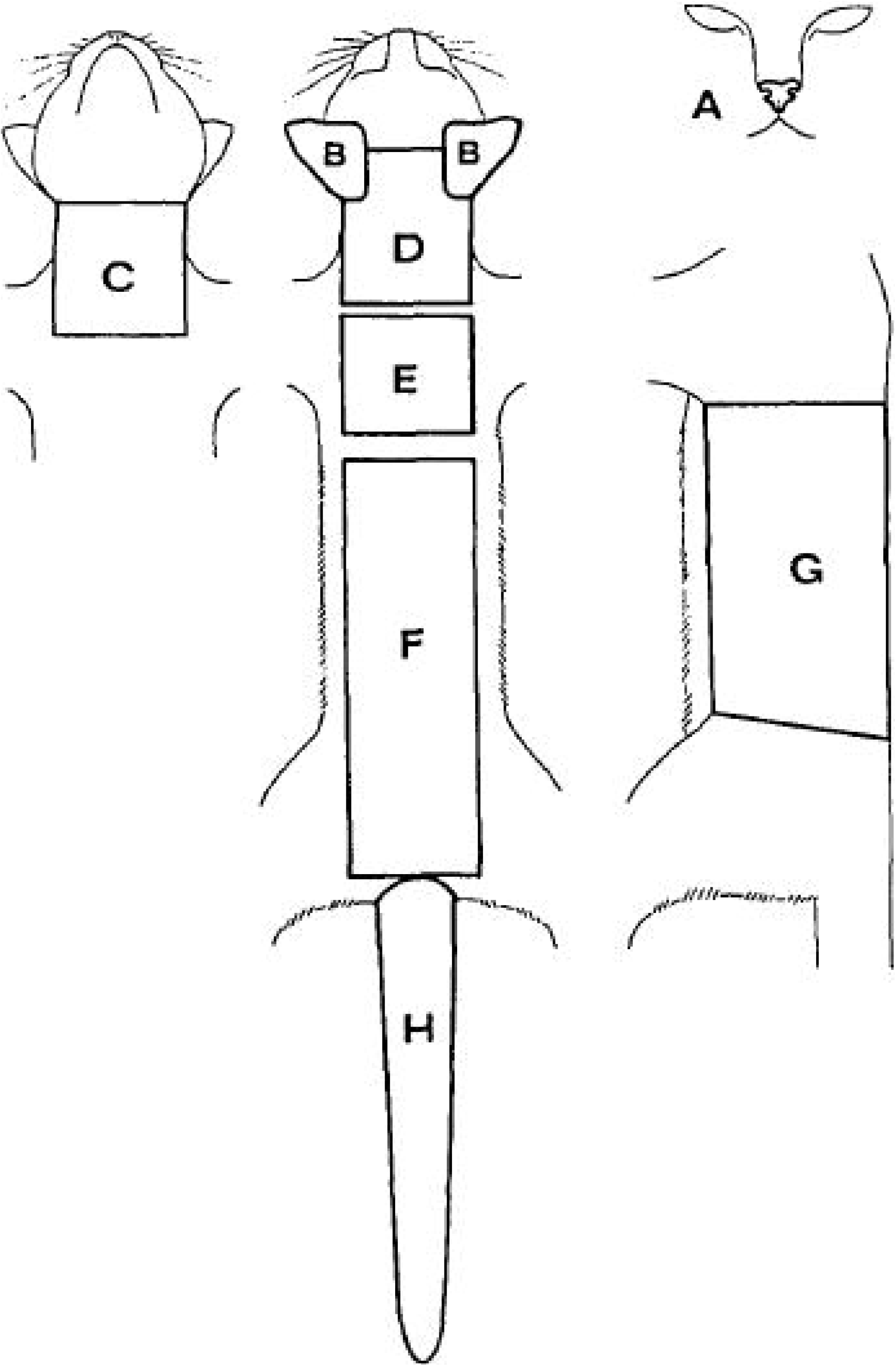
Somatic regions of *Felis silvestris*. A: *Rhinarium*, B: *Pinnae*, C: *Gularis*, D: *Occipitalis-Cervicalis*, E: *Scapularis*, F: *Dorsalis*, G: *Lateralis*, H: *Caudalis* (after Ragni and Possenti 1996).

**Fig. 3.**
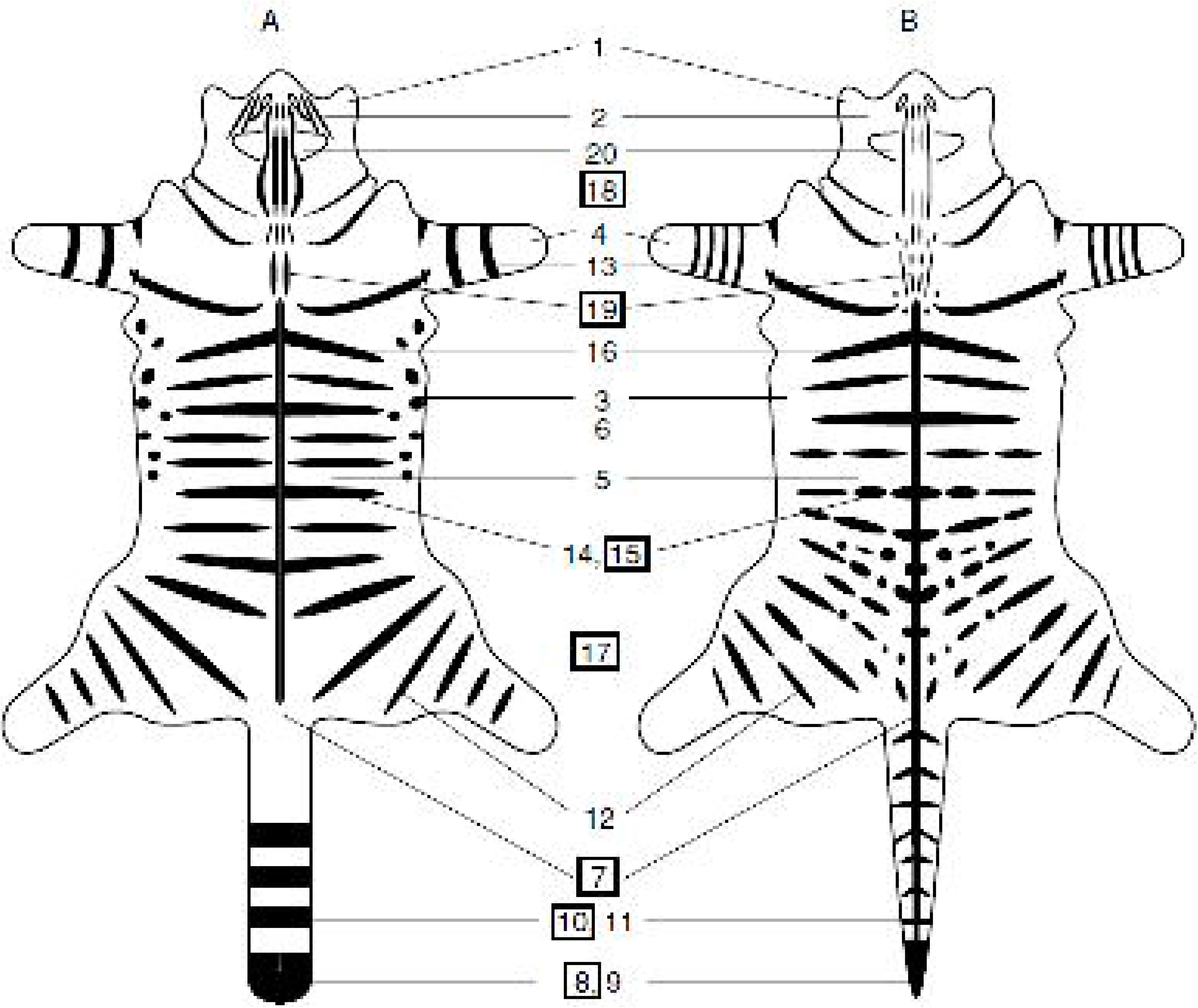
Comparison of pelage characters in Scottish wildcats (A) and putative domestic cats (B). 1: White on chin, 2: Stripes on cheek, 3: Dark spots underside, 4: White on paw, 5: White on flank, 6: White on back, <u>7: Extent of dorsal line, 8: Shape of tail tip</u>, 9: Colour of tail tip, <u>10: Distinctness of tail bands</u>, 11: Alignment of tail bands, 12: Stripes on hind leg, 13: Bands encircling foreleg, 14: Tabby coat patterns, 15: Broken stripes on flanks & hindquarters, 16: Stripes on body, <u>17: Spots on flanks & hindquarters, 18: Stripes on nape, 19: Stripes on shoulder</u>, 20: Colour of the back of ear (adapted from Kitchener et al. 2005). Properties best suited for discrimination of A and B are marked with black frames.

Despite these in-depth classification tools and several studies tackling differences in pelage characters between feline taxa, a systematic analysis of a possible link between pelage colour variations and the European distribution of *F. silvestris silvestris* has not been reported. Therefore, based on the pelage characteristics discussed by Ragni and Possenti (1996, Fig. 2) and Kitchener et al. (2005, Fig. 3) we compared 98 *F. silvestris silvestris* individuals from six distinct European areas (Fig. 1) by combining several cluster analysis methods to construct a plausible view on the relationship between feline coat patterning and geographic distribution. The rationale for the study is the hypothesis that clear differences in the fur patterning between the different regions exist.

## Material and methods

Based on the pelage character classification systems shown in Fig. 2 and Fig. 3 we investigated the pelage characteristics of 98 *F. silvestris silvestris* individuals from six European regions (21 from the Eifel, 49 from the Harz, 12 from the Caucasus, 4 from Western Spain, 10 from Switzerland or the North-East of France and 2 from the South-West of Greece). All individuals were adults, and cases in which the individual might have been a hybrid between wild- and domestic cats were excluded. Furthermore, sex was not taken into account. The specimens were studied at the following Institutions: Zentralmagazin Naturwissenschaftlicher Sammlungen der Martin-Luther-Universität Halle-Wittenberg, Zoologisches Forschungsmuseum Alexander König, **ZFMK**, Zoologisches Museum Berlin, **ZMB**, in Germany, Zoological Museum, University of Moscow, Moscow, Russia, **ZMNM**, and Museum d’Histoire Naturelle, Geneva, Switzerland, **MHNG**.

Polychoric correlations, as a binary version of Spearman’s rank correlation coefficient, were used as a measure of correlation between pelage characteristics. Pairwise tetra-choric (i.e. the special case of poly-choric for binary variables) correlations were computed with R 3.1.2. (R Core Team 2014a) using the external packages ‘polycor’ (Fox 2010) and ‘foreign’ (R Core Team 2014b). Cluster analyses were conducted with the commercial software SPSS 14.

A descriptive statistical analysis was also performed with SPSS 14 to determine the frequency of variables. Two types of analyses were made. The first analysis was performed on all categorical variables and highlighted variables unhelpful for cluster computation (i.e. variables with equal characteristic values for all individuals). Based on this and on general consideration, three pelage characteristics were discarded: (1) shape of tail tip (Fig. 3, characteristic 8) since only one individual (HA 89/154) had a pointed tail. HA 89/154 could not securely be identified as a wildcat and was therefore excluded from the analysis. All other individuals had wide tail tips, rendering the variable useless. (2) Basic lateral colouration and (3) basic thorax and abdomen colouration were also discarded since no standardised coding system for this character is available as yet.

For the analysis, variables were transformed from categorical variables into binary (dummy-) variables. A variable transformation was carried out with SPSS Syntax (see Appendix A for a SPSS Syntax code example). At this point, further variables could be discarded since all individuals had equal characteristic values of 0, i.e. absence of character (scap5 coding for irregular *Scapularis* pattern, fll4 coding for four bands encircling the foreleg and rhic3 coding for *Rhinarium* colours different from red and dark).

The final SPSS worksheet (a *.sav file) was imported to R using the ‘foreign’ package. With R, using the ‘polycor’ package, pairwise tetra-choric correlations between all binary variables were calculated and listed in a correlation matrix (see Appendix B for the R code used). To reduce the number of variables, one of two variables with correlations higher than 0.7 was discarded. Preferably, variables, such as the colouration of tail/back and chest spot were kept while variables with imprecise coding, such as paw characters or differentiation between clear and vague pelage spotting, were discarded. The initial amount of 89 dummy variables could be reduced to 22 uncorrelated binary variables. It is noteworthy that even though most categorical variables have more than two characteristic values and thus are not binary, each variable of our data can sufficiently be explained by precisely one binary variable. Additionally, one of the three different tail characteristics (tail for the number of tail-circling dark bands around the tail) can be discarded since it is explained by the remaining binary variables (Tab. 1).

**Table 1.** Comparison of the initial set of categorical variables and the set of binary variables used for cluster analysis. The set of binary variables was obtained by first calculating dummy variables from the full set of categorical variables and follow-up analysis of tetrachoric correlation between the binary variables. From two variables with correlation higher than 0.7 one variable was discarded (see text). The number of characteristic values for categorical variables is given in brackets. For binary variables the number of characteristic values naturally is two

| Full set of categorical variables | Reduced binary variable set |
| --- | --- |
| rhic (2) | rhic1 |
| rhir (2) | rhir1 |
| lip (3) | lip1 |
| che (2) | che1 |
| pinp (2) | pinp1 |
| pinc (2) | pinc1 |
| gulb (4) | gulb2 |
| gulw (4) | gulw1 |
| fore (2) | fore1 |
| occp (4) | occp1 |
| occl (5) | occl1 |
| scap (4) | scap1 |
| dors (10) | dors1 |
| latp (5) | latp1 |
| taila (2) | taila1 |
| tailr (7) | tailr3 |
| tailu (6) | - |
| venp (3) | venp1 |
| dleg (3) | dleg1 |
| dpc (6) | dpc1 |
| vea (3) | vea1 |
| flp (4) | flp1 |
| fll (5) | fll1 |

The second descriptive statistical analysis was performed on the reduced set of binary variables to visualise character distribution among individuals. Cluster analysis was carried out with reduced characteristics sets (Tab. 2). Missing values were excluded list by list, giving invalid individuals. Two types of cluster analyses, the hierarchical and the two-step cluster analysis, were made with different specifications in terms of variable choice and distance measures as well as clustering methods (Tab. 2).

**Table 2.**
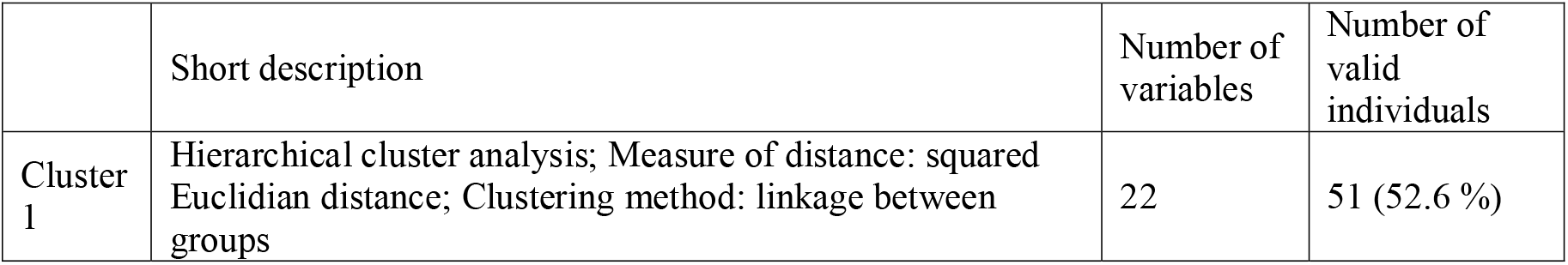
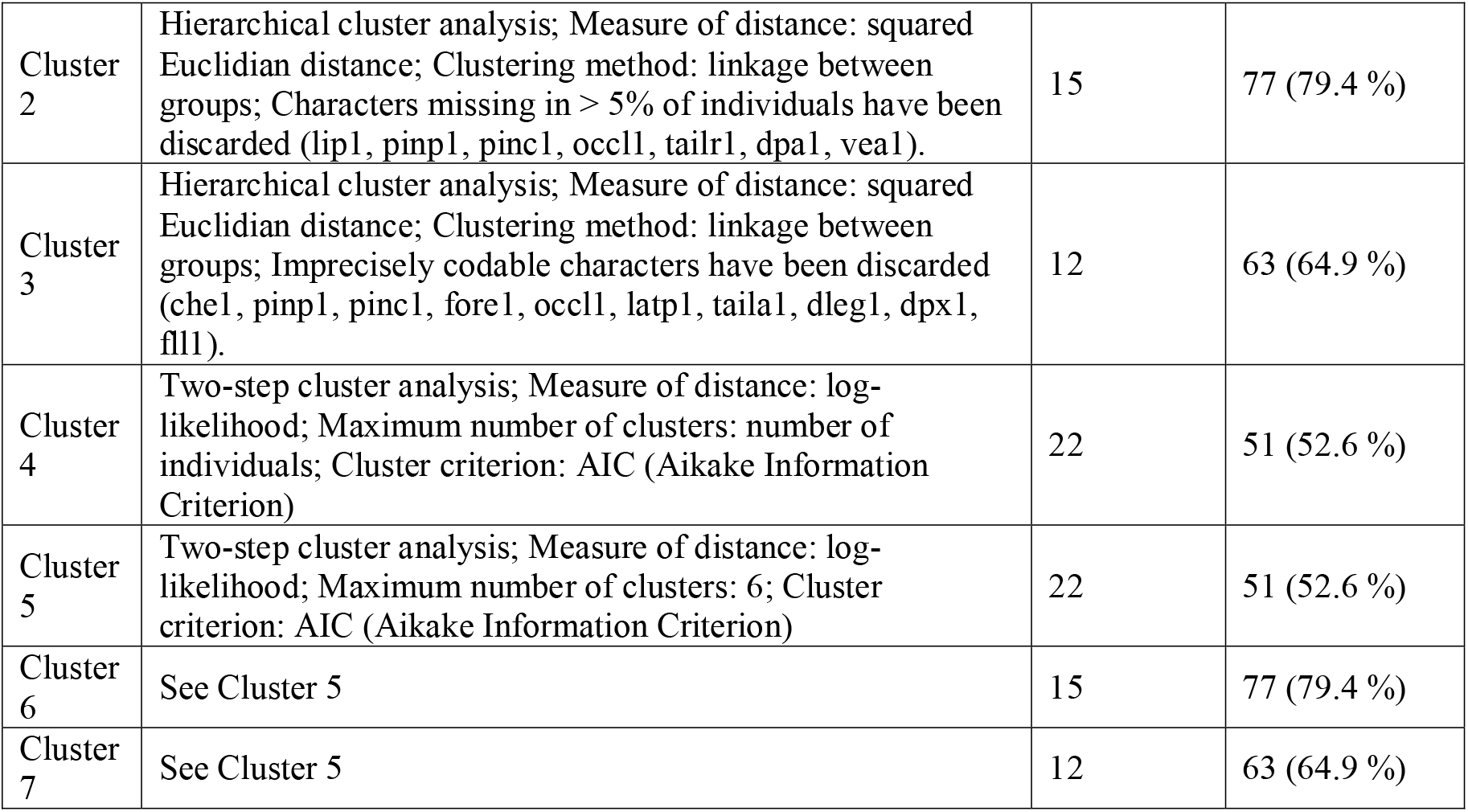
Summary of the seven conducted cluster analyses. Three analyses were carried out with a hierarchical cluster analysis, squared Euclidian distance as distance measure and linkage between groups as clustering method; the other four analyses were carried out with a two-step cluster analysis, in which hierarchical clustering was used for pre-clustering and as a fundament for clustering based on log-likelihood with AIC as final cluster criterion (see text). Further differentiation was due to number of analysed variables and predefined number of clusters

In hierarchical cluster analyses individuals bearing similar characteristics build clusters. As binary variables are metric, similarity could be measured by squared Euclidian distance. Distances between built clusters were measured as distance between cluster averages (i.e. average linkage between groups). Consequently, clusters were built iterative in an agglomerative bottom-up approach from a pre-calculated distance matrix. The results were presented in dendrograms.

Two-step cluster analyses, as the term suggests, derive final clusters from groups formed in a pre-clustering process. Pre-clustering is similar to hierarchical clustering. The obtained groups, also clusters, are then assessed by the product of single maximised log-likelihood functions for every variable, assuming a binominal distribution of the binary variables. The maximum likelihood approach gives the most probable and plausible clustering based on the number of final clusters given by the user. We set this input to the expected value of six clusters as individuals from six geographic regions were compared. Then, the maximum likelihood model for six clusters is calculated, i.e. the individuals are grouped into six clusters so that the product of the six single log-likelihood functions is maximal. If no number of final clusters is specified by the user (Cluster 4), *n* maximum likelihood models are calculated with *n* being the number of valid individuals used for clustering. For each of these maximum likelihood models, the respective predefined number of clusters is set to *n*. After that, the Akaike information criterion (AIC) is calculated for each maximum likelihood model. The AIC is a criterion that optimises model selection by a trade-off between predictability measured by the weighted least square method and simplicity and defined by the number of degrees of freedom minus number of parameters as a type of ‘Occam’s razor’ parsimony principle. Consequently, the model with the lowest AIC value is selected as the most plausible (see also description of cluster analyses in Brosius, 2006).

Two-step cluster analyses do not output dendrograms but add cluster variables to the data sheet. To visualise the individuals’ cluster affiliation, a dummy hierarchical cluster analysis was conducted in which only the cluster variable was taken into account. This leads to non-hierarchical dendrograms with equal distances between clusters (Cluster 4 to 7, Fig. 12 to 18).

Both types of cluster analyses were performed with different sets of variables (full, reduced by characters absent in more than 5 % of the individuals and reduced by imprecisely codable variables). Reduced variable sets increased the number of valid individuals.

Resulting clusters were categorised into non-closed and closed clusters, the latter being clusters which contain all valid individuals from one region and no valid individuals from any other region.

## Results

### Descriptive Statistics

Based on the first descriptive analysis, no characteristics could be discarded as all characteristic values were found in at least one individual. Consequently, all variables could be used to differentiate individuals. The most one-sided character is che (stripes on cheek), for which 88.5 % of all valid individuals (85 out of 96) were found to have the same characteristic value, i.e. two clear stripes (note that che already is a binary variable before variable transformation).

In the second descriptive statistical analysis che (i.e. che1) remained the most one-sided variable followed by fll1 (one band encircling the foreleg) which is present in 88.2 % (82 out of 93) valid individuals (not shown). All other variables showed more balanced distributions.

### Cluster analysis

Figures 12 to 18 (Appendix C) show the respective results of the cluster analyses presented in Tab. 2 as full dendrograms (Cluster 1 to Cluster 7). Here, we highlight excerpts from those dendrograms. Cluster 4, the only cluster analysis in which the number of clusters was not predefined, produced 7 clusters.

The Eifel region (region 1, initial n=21) was not represented in a closed cluster for any of the cluster analyses. Rather, individuals from that region can be found in many clusters (not shown). The same results were obtained for the Harz region (region 2, initial n=49) although all analyses except Cluster 6 produced some clusters which contained individuals from the Harz only. However, neither of the cluster analyses produced closed clusters for the Harz region (Fig. 4).

**Fig. 4.**
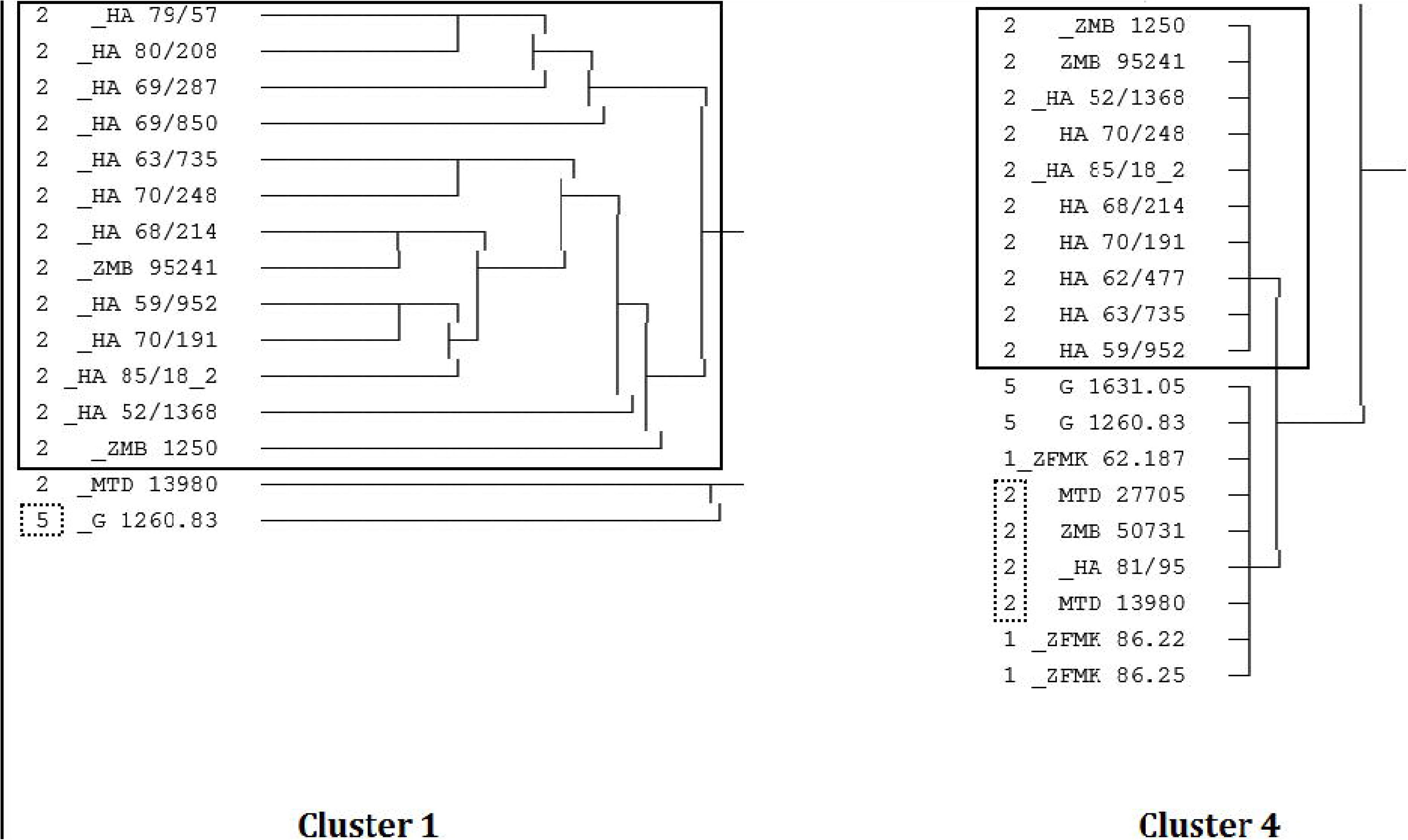
Exemplary comparison of results from Cluster 1 and Cluster 4 for the Harz region. Clusters with a frame only contain individuals from the Harz region. The dotted frame indicate individuals that prevent the cluster from being closed. For the Cluster 1 result the potentially closed cluster contains an individual from region 5 (Switzerland/North-Eastern France), while for the Cluster 4 result, not all individuals from region 2 were grouped together. Note that Cluster 4 did not output hierarchical clusters.

For the Caucasus (region 3, initial n=12), the most remote region in terms of distance from a hypothetical centroid among all regions, some analyses produced closed clusters (Cluster 4, note that the amount of clusters was not predefined in Cluster 4, and Cluster 5, Fig. 5).

**Fig. 5.**
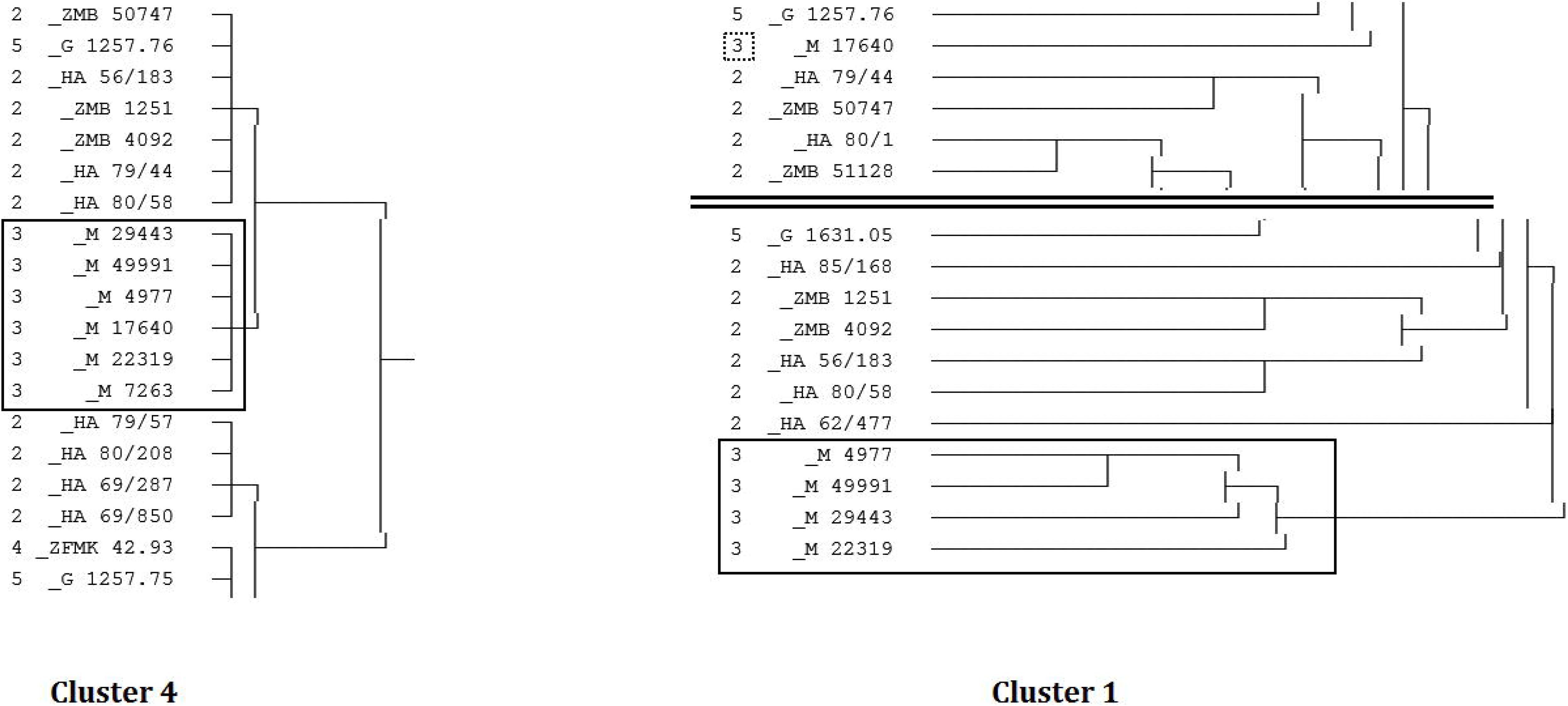
Results from Cluster 4 and Cluster 1 for the Caucasus region. Cluster 4: The cluster with a frame only contains individuals from the Caucasian region. The cluster is closed as no individual from this region was grouped into another cluster and no individual from other regions was grouped into the Caucasus cluster. Cluster 5 produced the same result. Note that Cluster 4 did not output hierarchical clusters. Cluster 1: The cluster with a frame only contains individuals from the Caucasian region. The individual with a dotted frame indicates an individual from the Caucasian region that was not grouped into the main cluster for that region. Therefore the cluster is not closed. However, an out-group is formed by the Caucasian cluster. The black double line indicates that in this image a part of the original dendrogram is missing.

Both these closed clusters differed from all other clusters in two characteristics, i.e. the number of stripes on the *Occipitalis-Cervicalis* (with the binary variable occl1 coding for five or more stripes) and the pelage patterning on the *Lateralis* (with the binary variable latp1 coding for a disorderly and blurred patterning). Our results indicate that wildcats with five or more stripes on the *Occipitalis-Cervicalis* tend to be Caucasian wildcats (Tab. 3 for Cluster 4) as 83.3 % of all valid individuals with five or more stripes were Caucasian while only 4.4 % of non-Caucasian individuals showed this phenotype. Even though these stripes were uncontinuous and conjunct in only few individuals, i.e. 19.6 % (10 out of 51), they always were continuous and separate in Caucasian wildcats (not shown). A similar but less explicit distribution can be observed for the *Lateralis* patterning: wildcats with a clear patterning with little blurriness tend to be Caucasian (not shown).

**Table 3.** Exemplary characteristic value distribution for the occl1 variable in analysis Cluster 4. The calculated cluster 7 is a closed cluster with Caucasian individuals only. 71.4 % of all valid individuals with five or more stripes on the Occipitalis-Cervicalis come from the Caucasus region. Note that 83.3 % (5 out of 6) Caucasian cats have five or more stripes while only 4.4 % (2 out of 45) non-Caucasian cats have five or more stripes. O indicates absence, 1 presence of a character

|  | 0 |  | 1 |  |
| --- | --- | --- | --- | --- |
| Cluster | Frequency | Percentage | Frequency | Percentage |
| 1 | 10 | 22.7 % | 0 | 0 % |
| 2 | 8 | 18.2 % | 1 | 14.3 % |
| 3 | 8 | 18.2 % | 0 | 0 % |
| 4 | 4 | 9.1 % | 0 | 0 % |
| 5 | 6 | 13.6 % | 1 | 14.3 % |
| 6 | 7 | 15.9 % | 0 | 0 % |
| 7 | 1 | 2.3 % | 5 | 71.4 % |
| Combined | 4 | 100 % | 7 | 100 % |

Cluster 1 and Cluster 3 still yielded merely Caucasian clusters with 4 individuals each. However, these clusters were not closed since several individuals were integrated into other clusters. It is noteworthy that the Caucasian cluster in Cluster 1 is, compared to all other clusters, the outgroup, i.e. the Caucasian individuals in that cluster have the highest distance from all other individuals (Fig. 5). In Cluster 6 and 7 Caucasian individuals were mainly grouped into one cluster, however among individuals from other regions. Furthermore, one (Cluster 7) or two (Cluster 6) Caucasian individual(s) were integrated into other clusters (Fig. 6). In Cluster 2 the scattering of Caucasian individuals was much more pronounced, even though three individuals were still grouped in proximity (Fig. 6).

**Fig. 6.**
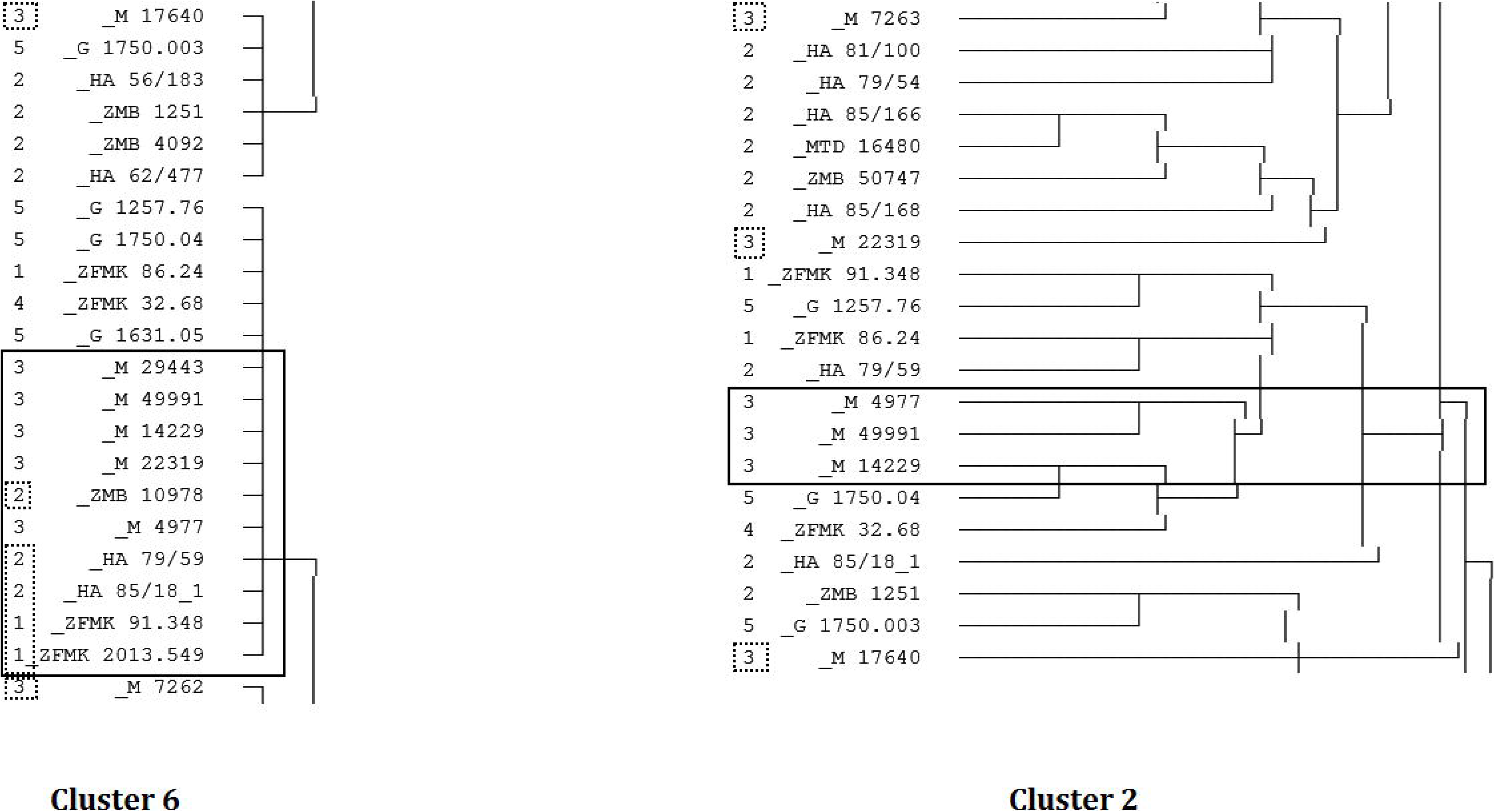
Results from Cluster 6 and Cluster 2 for the Caucasus region. The clusters with a frame mainly contain individuals from the Caucasian region. In Cluster 6 some individuals from other regions have been grouped into the Caucasian cluster, as well as some Caucasian individuals were not grouped into the main cluster for that region and in Cluster 2 one Caucasian individual was grouped outside the main cluster (dotted frame). Therefore the clusters are not closed. Note that Cluster 6 did not output hierarchical clusters.

Two individuals (ZFMK 42.84 and ZFMK 42.93) from Western Spain (region 4, initial n=4) were ever-present and always grouped into the same cluster (closed clusters in Cluster 1, 3). However, in Cluster 2 the cluster is not closed since another Spanish individual was grouped into a different cluster (Fig. 7).

**Fig. 7.**
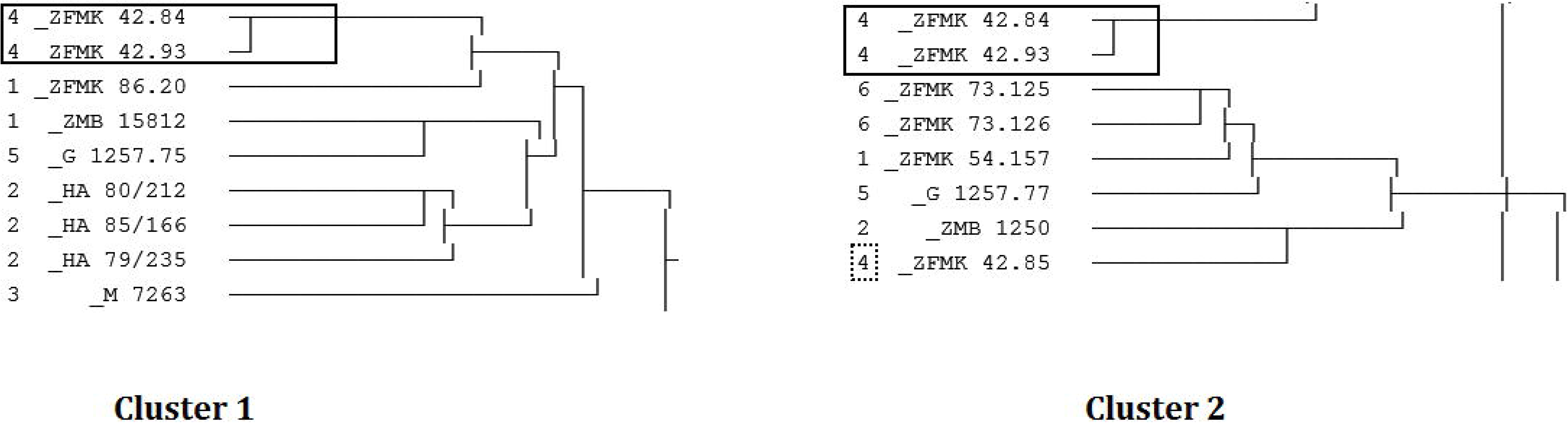
Results from Cluster 1 and Cluster 2 for the Western Spain region. The clusters with a frame only contain individuals from the Spain region. For Cluster 1 the cluster is closed while for Cluster 2 another individual from Western Spain was grouped outside the cluster (dotted frame). Therefore the cluster is not closed.

In the remaining analyses (Cluster 4 to 7, note that all of these were two-step cluster analyses without hierarchical clustering) ZFMK 42.84 and ZFMK 42.93 were grouped into the same cluster among individuals from various other regions (Fig. 8).

**Fig. 8.**
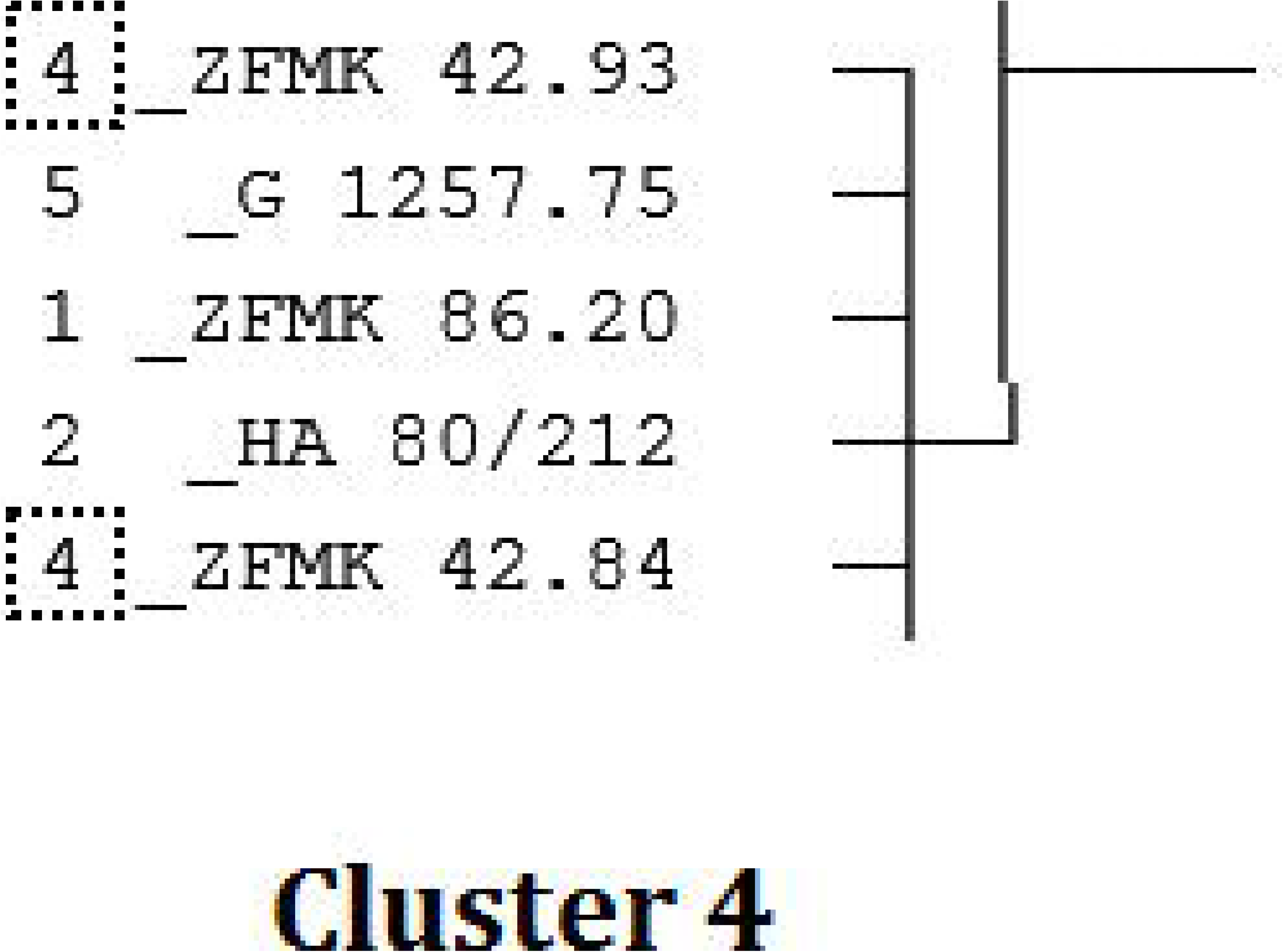
Exemplary result from Cluster 4 for the Western Spain region. The two valid individuals from Western Spain are grouped into the same cluster (dotted frame), however, among several individuals from various other regions. Therefore the cluster is not closed. Note that Cluster 4 did not output hierarchical clusters.

Individuals from Switzerland/North-Eastern France (region 5, initial n=10) were always highly scattered over the dendrograms (not shown).

Individuals from Greece (region 6, initial n=2) were often discarded due to missing values. However, if present, the two individuals were grouped together (closed cluster in Cluster 2 and among other individuals from various other regions in Cluster 6, Fig. 9).

**Fig. 9.**
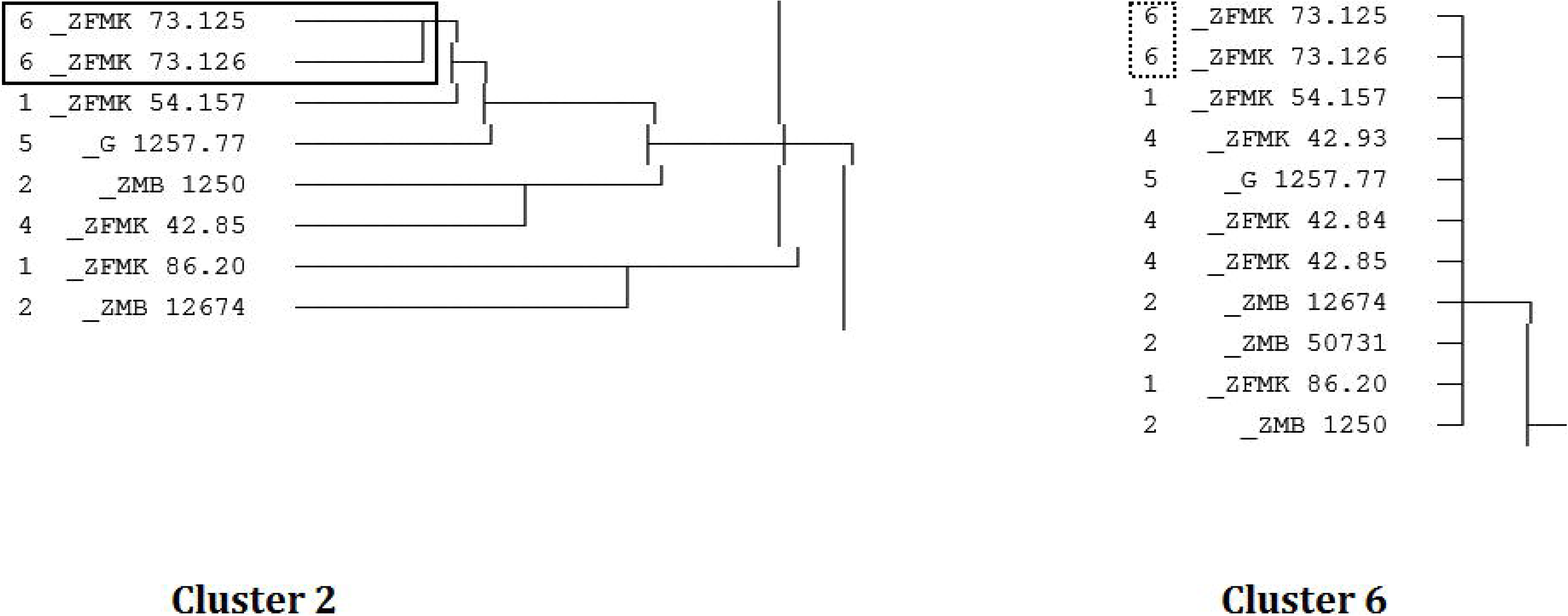
Results from Cluster 2 and Cluster 6 for the South-Western Greece region. The two individuals from South-Western Greece were grouped into the same cluster if present in the analysis. In Cluster 2, the cluster is closed (frame) while in Cluster 6 the Greek individuals were grouped among individuals from various other regions (dotted frame). Therefore the cluster is not closed. Note that Cluster 6 did not output hierarchical clusters.

## Discussion

A strong interpretation of our hypothesis that *F. silvestris silvestris* wildcats from different European regions can be distinguished by a coat pattern analysis, results in the expectation that all valid individuals can be grouped into six closed clusters. The predefined cluster number in two-step cluster analyses therefore has been six. However, our results do not show a clear correlation between combinations of pelage characteristics and geographic distribution, which is epitomised by the fact that Cluster 4, the only analysis without a predefinition, produced seven clusters. Consequently, the hypothesis needs to be rejected based on our data.

This is also supported by our findings for individuals from either the Eifel region (region 1) or Switzerland/North-Eastern France (region 5) that were dispersed over various clusters in all analyses (not shown). The same holds true for individuals from the Harz region (region 2), which occur in almost every non-closed cluster, though to a lesser extent, as almost all analyses do have clusters containing individuals from the Harz region only (Fig. 4). However, 50% (49 out of 98) of all individuals incorporated in the analysis were sampled in the Harz region. Hence, it needs to be taken into account that the ample distribution of individuals from the Harz is rather a statistical artefact than due to pelage characteristics meaning that simply too many Harz individuals were analysed as compared to individuals from other regions. This may result at a higher ratio of clusters containing individuals from the Harz region only than expected.

As opposed to that, individuals from the South-West of Greece (region 6), Western Spain (region 4) and especially the Caucasus (region 3) were grouped into clusters that proved to be relatively robust over the different cluster analyses. Both South-West Greece and Western Spain were represented in closed clusters for a hierarchical cluster analysis each (Western Spain in Cluster 1, South-West Greece in Cluster 2). In two-step cluster analyses (Cluster 4 to 7) the clusters were not closed, possibly, since only few individuals, two from South-West Greece and four from Western Spain, were analysed. Here, it must be taken into account that in two-step analyses, either with or without a predefined number of clusters, no cluster hierarchy is calculated. In contrast to hierarchical analyses where clusters are built bottom-up, in two-step analyses no intra-cluster differentiation takes place.

Accordingly, cluster classification is less detailed as clusters remain on the first level of differentiation without further clustering of individuals within already built clusters. Furthermore, we found rather uniform cluster sizes for the two step analyses. This means that clusters of two or four individuals did not occur. Hence, individuals from regions where only small sample sizes were collected, i.e. South-West Greece and Western Spain, are less likely to occur in clusters without individuals from any other region.

The strongest sign of coat differences from other European wildcats was found in Caucasian wildcats which were represented in closed clusters for two different analyses. Most strikingly, Cluster 4 – without a predefined number of clusters – and Cluster 5, differing from Cluster 4 only in the predefined number of six clusters, produced very similar results (Fig. 5). The two closed clusters found in these two analyses differed from all other clusters in distinct characters (Tab. 3). Especially the number of stripes on the *Occipitalis-Cervicalis* may be useful to differentiate between Caucasian and non-Caucasian wildcats as 83.3 % of all Caucasian wildcats have five or more stripes on the *Occipitalis-Cervicalis* while only 4.4 % of non-Caucasian wildcats show this phenotype. Furthermore, these stripes were separate and continuous in all Caucasian individuals while about one-fifth of all valid individuals also showed conjunct or uncontinuous stripes (not shown).

It is generally plausible that individuals from the Caucasian rather than any other region vary from all other individuals if, for any of the investigated regions, closed clusters would be expected for the Caucasus region more than for any other region, due to its comparatively high geographic separation which might be reflected in morphological separation. This separation is slightly supported by the fact that in the dendrogram calculated in Cluster 1 four Caucasian individuals form an outgroup (Fig. 5). This means that all valid individuals from this analysis are more similar to one another than to these four Caucasian individuals. However, two Caucasian individuals were found outside of the out-group, meaning they are less similar to other Caucasian wildcats than they are to wildcats from non-Caucasian regions (Fig. 5). As a consequence of this, our data suggest that there may be differences between the pelage characteristics of Caucasian and non-Caucasian wildcats, even though these may not be sufficient to clearly identify Caucasian individuals by pelage characteristics alone. This difficulty is illustrated by the fact that even though in all cluster analyses groups of Caucasian individuals were built, individuals from other regions were grouped among them in some analyses (Fig. 6).

In addition to the special concerns already mentioned, further general objections against hasty conclusions based on our results need to be made. As already implied, the sample size used in our work was small. Not only that our overall individual count of 98 is low, we also had to work with strongly varying sample sizes for each geographic region (from two individuals from South- Western Greece to 49 individuals from the Harz). But material available in collections of the other regions is rare or beyond reach within the scope of the current study. To account for this problem, further analyses with (i) a generally higher number of individuals and (ii) a more balanced selection of individuals from all regions need to be made. For now, our results can be interpreted as a tendency that if morphological differences exist between geographically separated populations within the *F. silvestris silvestris* species, these will likely be found between Caucasian and non-Caucasian wildcats or maybe also between wildcats from Western Spain and South-West Greece and other European regions.

## Conclusion

As our results do not show a clear relationship between the pelage characteristics and the geographic distribution of six distinct European *F. silvestris silvestris* populations, our hypothesis needs to be rejected. However, based on our data, a new hypothesis of coat patterning of Caucasian wildcats to some degree differing from other European wildcats may be constructed. Our results, however, do not provide hard evidence for this hypothesis as Caucasian wildcats were not grouped into closed clusters for all cluster analyses. To a lesser extent, pelage characteristic differences may also exist between wildcats from Western Spain as well as South-Western Greece and other European regions, although sample sizes from these two regions were very small. To test these refined hypotheses, more samples from the Caucasus and especially from Western Spain and South-Western Greece (but also Switzerland/North-Eastern France) need to be collected and compared according to our protocol.

## Acknowledgements

We would like to thank the following curators and staff of the collections under their care for the access and possibility to study the specimens: Dr. F. Mayr, Mrs. S. Jancke and N. Lnage, Berlin, Dr. K. Schneider, Halle, Dr. R. Hutterer, Bonn, Dr. V. Lebedev, Moskau and D. M. Ruedi in Geneva. Mrs. C. Schuster, Dresden, helped with the study of material in Halle and Berlin. Dr. M. Rudolf, Technical University Dresden commented on the statistics of the manuscript, Mrs. E. Orrison, Dresden, a native speaker, checked the English. The helpful comments of the editors and reviewers are gratefully acknowledged.

